# A leukemia-derived ENL/AF9 chemical probe enhances neuronal stress resilience and ameliorates ALS phenotypes

**DOI:** 10.64898/2026.04.09.717610

**Authors:** Luca Lo Piccolo, Chansunee Panto, Ranchana Yeewa, Ruedeemars Yubolphan, Saranyapin Potikanond, Salinee Jantrapirom

**Affiliations:** Center of Multidisciplinary Technology for Advanced Medicine (CMUTEAM), Faculty of Medicine, Chiang Mai University, Chiang Mai, Thailand; Drosophila Centre for Human Diseases and Drug Discovery (DHD), Faculty of Medicine, Chiang Mai University, Chiang Mai, Thailand; Department of Biochemistry, Faculty of Medicine, Chiang Mai University, Chiang Mai, Thailand 50200; Department of Pharmacology, Faculty of Medicine, Chiang Mai University, Chiang Mai, Thailand

**Keywords:** ENL/AF9, chromatin reader, chemical probe, integrated stress response, amyotrophic lateral sclerosis

## Abstract

Chemical perturbation of chromatin reader proteins provides a precise strategy to interrogate epigenetic control of neuronal stress adaptation. ENL and AF9 are YEATS-domain acyl-lysine readers best characterized in leukemia, but their roles in neurons remain unclear. Here, we use the selective YEATS inhibitor SR-0813 to define ENL/AF9 function in neuronal stress responses across *Drosophila* and human systems. SR-0813 phenocopies genetic ENL/AF9 reduction by extending lifespan and enhancing stress tolerance *in vivo*, and improves survival of human neurons under multiple stress conditions, with the strongest effects during endoplasmic reticulum stress. Mechanistically, SR-0813 attenuates PERK-ISR signaling and reduces apoptotic commitment without broadly enhancing proteostasis capacity. Notably, its effects are highly context dependent, conferring protection in stress-signaling-driven models but reduced efficacy or detrimental outcomes under chronic aggregation or mitochondrial stress. These findings identify ENL/AF9 as modulators of stress-response dynamics and highlight YEATS-domain inhibition as a context-dependent strategy to reshape neuronal resilience.

**GRAPHICAL ABSTRACT:** 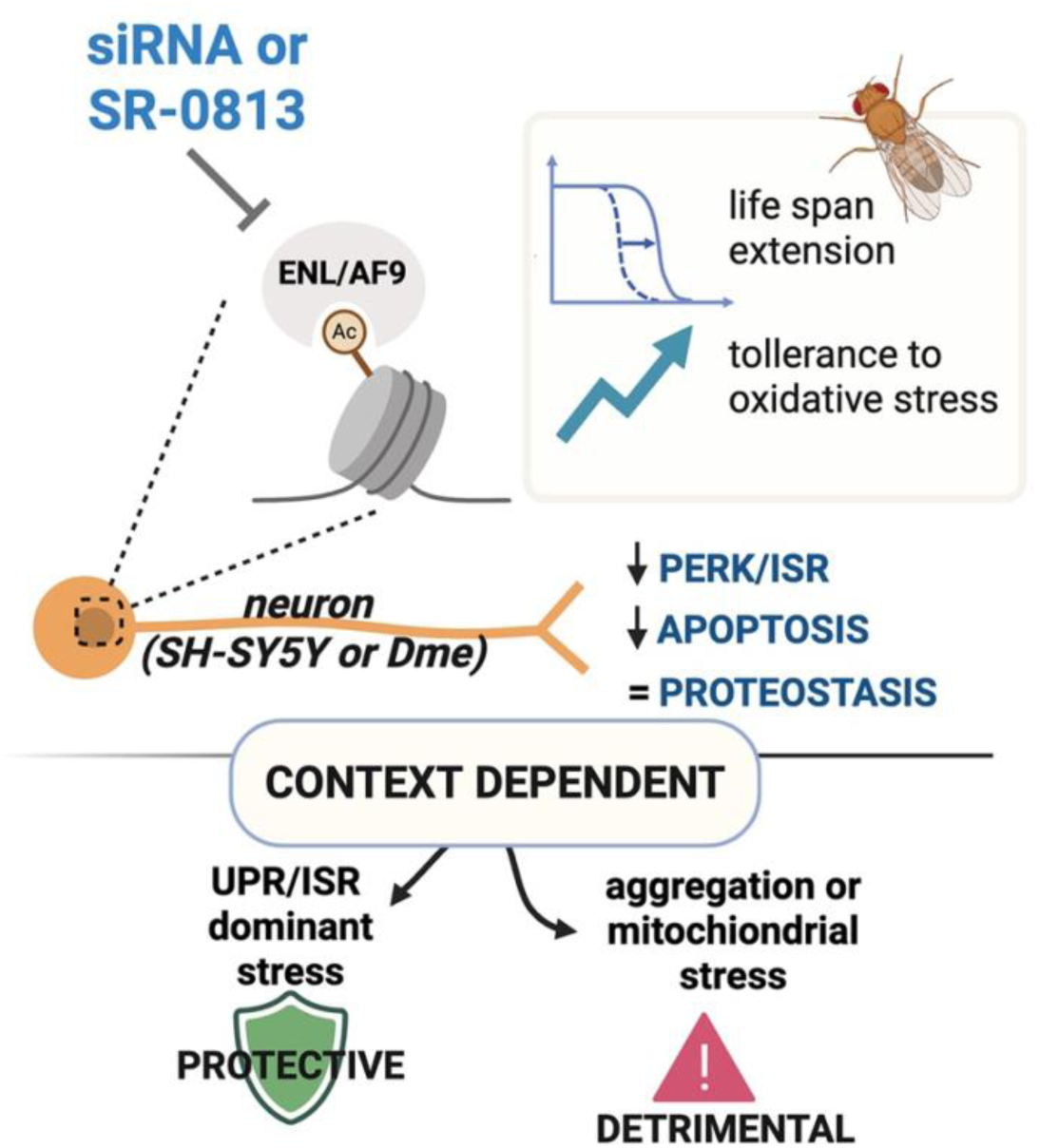

*siRNA* or the YEATS-domain inhibitor SR-0813 suppress the acyl-lysine reader ENL/AF9, extending lifespan and increasing H₂O₂ tolerance in *Drosophila*. In neurons (*Drosophila* and SH-SY5Y), ENL/AF9 inhibition lowers PERK–ISR signaling and apoptosis while maintaining proteostasis. Effects are context dependent: protective under UPR/ISR-dominant stress but potentially detrimental under aggregation or mitochondrial stress. Created with BioRender.

## INTRODUCTION

Neuronal identity and function rely on transcriptional programs that must be both stably maintained over long timescales and dynamically responsive to physiological and pathological stimuli. Chromatin-based regulation is now well established as a key determinant of neuronal development and plasticity, and as an important layer controlling stress-responsive transcriptional programs that shape neuronal resilience and vulnerability ^1–4^. However, the specific chromatin reader modules that tune these stress programs and their tractability to selective chemical perturbation remain incompletely defined.

One class of candidates with the potential to fulfill this role is the YEATS-domain family of chromatin readers ^5^. YEATS-domain proteins have emerged as selective sensors of histone acyl-lysine modifications, linking chromatin state to transcriptional output ^6^. ENL (MLLT1) and its paralog AF9 (MLLT3) function within transcriptional elongation complexes, including the super elongation complex (SEC), where YEATS-domain-dependent chromatin engagement promotes RNA polymerase II pause release and amplifies stimulus-responsive gene expression ^7–9^. Because histone acylation is tightly coupled to cellular metabolism through acyl-CoA availability, YEATS-domain recognition provides a potential mechanism by which metabolic state can be translated into transcriptional responses ^10,11^. While these features have been extensively studied in proliferative systems, particularly in cancer, whether YEATS-domain proteins regulate stress adaptation in post-mitotic neurons remains largely unexplored.

To begin addressing this question *in vivo*, we turned to *Drosophila melanogaster*, which provides a genetically tractable system with strong conservation of chromatin regulatory mechanisms ^12^. Flies encode a single ENL/AF9 ortholog, ear (here referred to as dENL/AF9), which contains a conserved YEATS domain and participates in related chromatin-associated complexes ^13–15^. We previously found that genetic reduction of *dENL/AF9* extends lifespan and enhances tolerance to oxidative stress, pointing to a role in organismal resilience ^16^. Notably, these effects are not shared by other YEATS-domain proteins, as perturbation of the *Drosophila* YEATS2 homolog (also known as D12) produces distinct neuronal phenotypes ^17^, indicating functional specificity within this protein family. Together, these observations raise the possibility that dENL/AF9 regulates neuronal stress responses through YEATS-dependent transcriptional mechanisms.

These findings prompted two key questions: whether the resilience phenotypes associated with *dENL/AF9* depletion are driven by reduced YEATS-domain activity, and whether this function is conserved in mammalian neuronal systems. To address this, we turned to SR-0813, a selective small-molecule inhibitor of the ENL/AF9 YEATS domains originally developed in the context of leukemia, where ENL-dependent transcription sustains oncogenic gene expression ^18^. Acting as a chromatin-competitive ligand, SR-0813 blocks YEATS-domain binding to acyl-lysine marks on histones, thereby disrupting ENL/AF9-dependent transcriptional programs while enabling acute and reversible interrogation of ENL/AF9 chromatin reader activity without broadly targeting other YEATS-domain-containing proteins such as YEATS2 or GAS41 ^19^. Although its activity toward dENL/AF9 has not been directly established, this conservation and functional similarity support the use of SR-0813 as a probe in this system when interpreted alongside genetic benchmarks. Here, we combine chemical perturbation with genetic approaches in *Drosophila* and human neuronal models to define how ENL/AF9 YEATS-domain function shapes neuronal stress adaptation across distinct physiological and disease-relevant contexts.

## RESULTS

### SR-0813 promotes longevity and oxidative stress resistance in *Drosophila* without intrinsic radical scavenging activity

To test whether pharmacological modulation of ENL/AF9 can recapitulate the beneficial effects observed upon genetic depletion, we administered the selective inhibitor SR-0813 to *Drosophila* and assessed phenotypes previously associated with dENL/AF9 reduction. We chose SR-0813 because it is the best-characterized, commercially accessible dual ENL/AF9 YEATS inhibitor, with high biochemical potency, strong selectivity over other YEATS domains, and a well-validated chromatin-competitive mechanism that evicts ENL from target loci without inducing protein degradation ^19^.

Adult flies were maintained on standard food until day 20 (D20) and subsequently transferred to food containing SR-0813 (1 or 10 μM), with regular media replacement (Figure 1A). Kaplan–Meier survival analysis revealed a significant extension of lifespan at both concentrations compared to untreated controls (Figure 1B), indicating that pharmacological inhibition phenocopies the longevity benefits of dENL/AF9 reduction.

**Figure 1.**
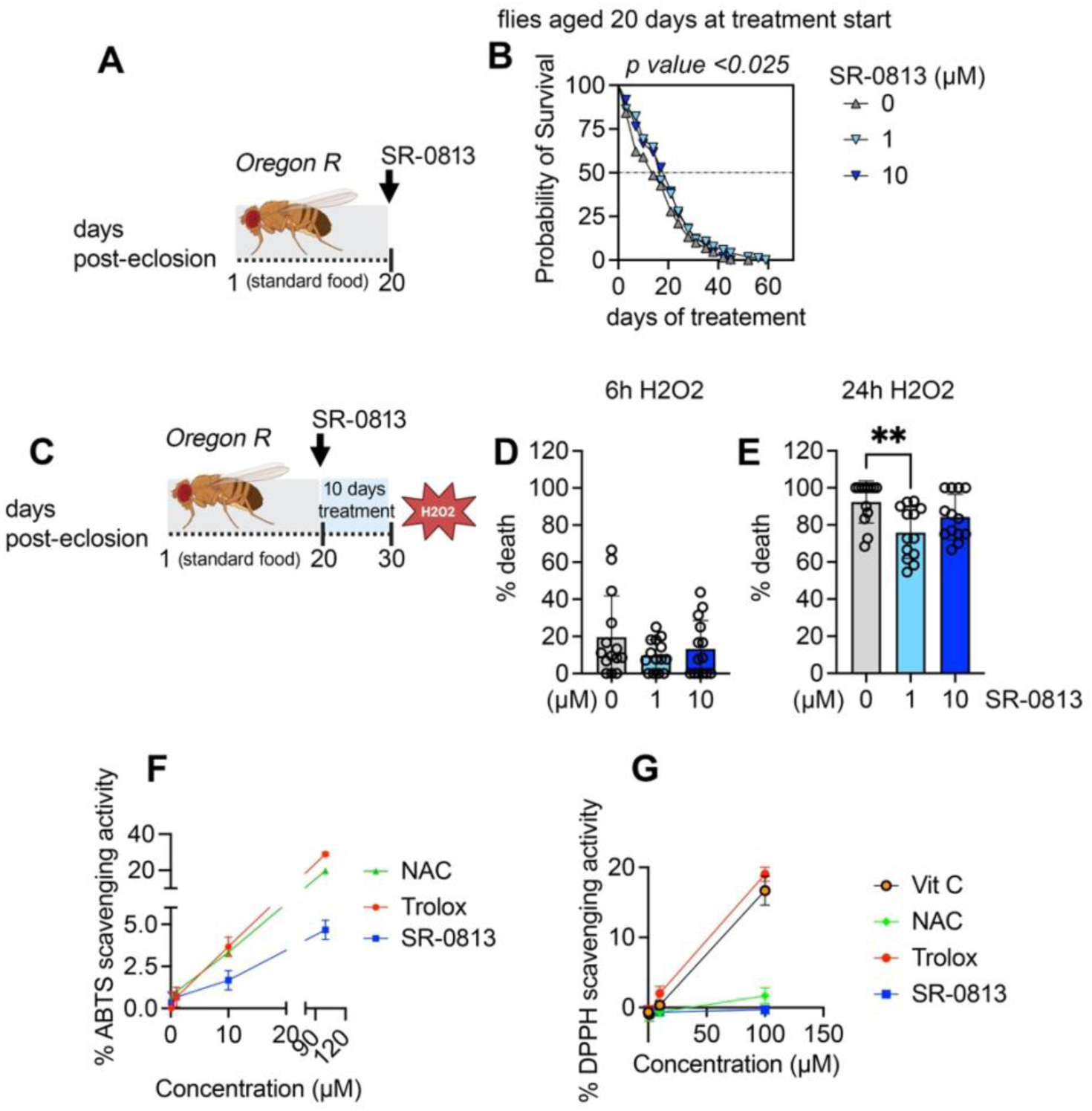
SR-0813 phenocopies genetic ENL/AF9 inhibition to enhance stress resistance in *Drosophila* without direct antioxidant activity. (A) Experimental paradigm for lifespan analysis. Oregon R flies were aged on standard food until day 20 (D20) and then transferred to food containing SR-0813 (1 or 10 μM) or vehicle control, and survival was monitored over time. (B) Kaplan–Meier survival curves of flies treated as described in (A). Statistical significance was determined using the Mantel–Cox test. (C) Experimental paradigm for oxidative stress assays. Flies were aged to D20, treated with SR-0813 (1 or 10 μM) for 10 days, and subsequently exposed to hydrogen peroxide (H_2_O_2_) 5% for 6 or 24 h before survival assessment. (D) Survival following 6 h (H_2_O_2_) exposure under the indicated conditions. (E) Survival following 24 h (H_2_O_2_) exposure under the indicated conditions. Individual values are shown with mean ± SD. (F–G) Radical scavenging activity of SR-0813 measured using (F) 2,2′-azino-bis(3-ethylbenzothiazoline-6-sulfonic acid (ABTS) or (G) 2,2-diphenyl-1-picrylhydrazyl (DPPH) assays across increasing concentrations, with vitamin C (vit C), Trolox, and N-acetylcysteine (NAC) as positive controls. For survival assays, n = 100 flies per condition. Statistical significance in (D) and (E) was determined using one-way ANOVA with appropriate multiple-comparisons testing. Data are presented as mean ± SD.

We next examined whether SR-0813 similarly enhances tolerance to oxidative stress. Following 10 days of treatment initiated at D20, flies were exposed to hydrogen peroxide (H₂O₂) (Figure 1C). While short-term exposure (6 h) resulted in minimal mortality with no detectable effect of treatment, prolonged exposure (24 h) led to substantial lethality that was partially rescued by SR-0813, particularly at the lower concentration (Figures 1D and 1E).

Because improved survival under oxidative stress can result from direct antioxidant activity, we evaluated the radical scavenging properties of SR-0813. In both ABTS and DPPH assays, SR-0813 did not exhibit detectable scavenging activity across the tested concentrations (Figures 1F and 1G), in contrast to established antioxidants.

Together, these findings show that SR-0813 recapitulates key phenotypes associated with genetic reduction of dENL/AF9, enhancing lifespan and stress tolerance through mechanisms unlikely to involve direct antioxidant effects. Given that SR-0813 selectively targets the ENL/AF9 YEATS domain, these results suggest that the lifespan extension and increased stress tolerance observed upon dENL/AF9 depletion are at least in part mediated by reduced YEATS-domain activity and consequent transcriptional reprogramming.

### SR-0813 enhances stress resilience in differentiated human neurons and attenuates apoptosis under ER and proteotoxic stress

The ability of SR-0813 to phenocopy neuronal ENL/AF9 depletion in *Drosophila* prompted us to examine whether its protective effects extend to human neuronal systems. To this end, we evaluated its impact in retinoic acid (RA)-differentiated SH-SY5Y cells, a widely used model of human neurons.

Differentiation with RA (10 μM, 3 days) established a neuronal-like phenotype suitable for downstream analyses (Figure S1). Under these conditions, treatment did not affect cell viability across the tested dose range (Figure S2), indicating a lack of intrinsic toxicity to neurons. To challenge neuronal resilience, we exposed differentiated cells to multiple stress paradigms relevant to neurodegeneration, including oxidative stress (H₂O₂), proteasomal inhibition (MG132), endoplasmic reticulum (ER) stress (tunicamycin, Tm), and mitochondrial dysfunction (rotenone) (Figure S3). Cells were co-treated with increasing concentrations of the compound during 48 h of stress exposure (Figure 2A).

**Figure 2.**
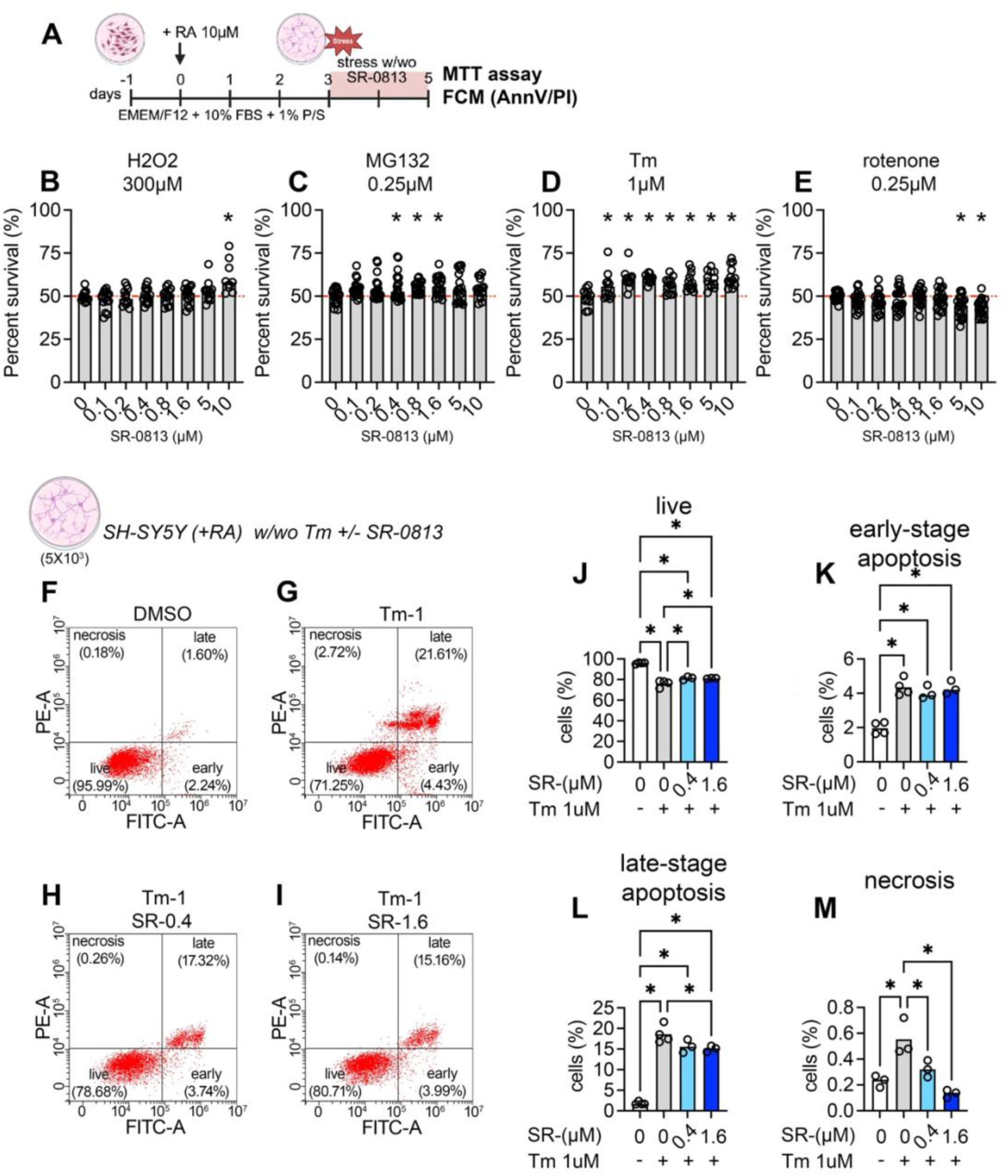
SR-0813 protects differentiated SH-SY5Y cells from ER and proteotoxic stress–induced loss of viability. (A) Experimental paradigm. SH-SY5Y cells were differentiated with retinoic acid (RA; 10 µM) for 3 days and then exposed for 2 days to H₂O₂ (300 µM), MG-132 (0.25 µM), tunicamycin (Tm; 1 µM), or rotenone (0.25 µM), with vehicle (DMSO) or SR-0813 (0–10 µM). (B–E) MTT viability assay of differentiated SH-SY5Y cells exposed to (B) H₂O₂, (C) MG-132, (D) Tm, or (E) rotenone in the presence of increasing SR-0813 concentrations. Each dot represents one technical replicate; data are from 3 independent experiments. (F–I) Representative Annexin V/propidium iodide (PI) flow cytometry plots from cells treated with (F) DMSO, (G) Tm, (H) Tm + SR-0813 (0.4 µM), or (I) Tm + SR-0813 (1.6 µM). (J–M) Quantification of Annexin V/PI populations showing (J) live cells (Annexin V⁻/PI⁻), (K) early apoptosis (Annexin V⁺/PI⁻), (L) late apoptosis (Annexin V⁺/PI⁺), and (M) necrosis (Annexin V⁻/PI⁺). Each dot represents one technical replicate; data are from 3 independent experiments. SR-indicates SR-0813. For (B–E) and (J–M), statistical significance was assessed by ordinary one-way ANOVA with Sidak’s multiple-comparisons test (p < 0.05).

Treatment enhanced neuronal survival under H₂O₂ (300 μM), MG132 (0.25 μM), and Tm (1 μM), while exacerbating toxicity under rotenone (0.25 μM) (Figures 2B–2E). The most pronounced protective effect was observed under ER stress, where increased survival was detected from low concentrations (0.1 μM) across the entire dose range tested. Under proteasomal stress, a peak protective effect was observed in the range of 0.4–1.6 μM, whereas under oxidative stress, significant benefit was limited to higher concentrations (10 μM) (Figures 2B–2E). Based on these profiles, subsequent analyses focused on Tm- and MG132-induced stress.

To assess whether the survival effects observed in the MTT assay reflect modulation of cell death pathways, we performed Annexin V/propidium iodide (PI) flow cytometry in differentiated SH-SY5Y cells, with single-stain controls used to define gating boundaries (Figures S4C–S4E).

Under ER stress, Tm induced marked apoptotic and necrotic cell death (Figures 2F–2G), accompanied by neurite disruption as observed by light microscopy (Figure S4B). Co-treatment increased the proportion of viable cells while reducing late-stage apoptosis and necrotic populations in a dose-dependent manner, with maximal protection observed at 1.6 μM (Figures 2J–2M and S4B).

We next applied the same approach to proteasomal stress. MG132 induced a more severe phenotype than Tm, with pronounced morphological disruption (Figure S5B). Co-treatment modestly increased the proportion of viable (Annexin V⁻/PI⁻) cells, with a reduction in early apoptotic populations observed primarily at the lower concentration (0.4 μM), while no clear effect was detected on late apoptosis or necrosis (Figure S5A). Thus, consistent with the viability data, the protective effect under proteasomal stress is associated with a limited attenuation of cell death, less pronounced than that observed under ER stress.

Together, these findings indicate that SR-0813 enhances neuronal stress resilience, most prominently under ER stress, in part through attenuation of apoptotic cell death, prompting further investigation into its effects on ER stress signaling pathways.

### SR-0813 attenuates ROS accumulation and dampens ER stress, ISR, and apoptotic signaling

Given the pronounced protective effect observed under ER stress conditions, we next sought to define the molecular pathways underlying this phenotype. Because tunicamycin-induced toxicity is closely linked to oxidative stress and activation of maladaptive unfolded protein response (UPR)/integrated stress response (ISR) signaling, we first asked whether SR-0813 modulates these processes.

We began by assessing intracellular reactive oxygen species (ROS), a key contributor to ER stress–associated toxicity. In undifferentiated SH-SY5Y cells, tunicamycin induced a marked increase in ROS levels, as detected by CM-H₂DCFDA confocal imaging, which was progressively reduced by co-treatment in a dose-dependent manner (Figures 3A–3C). To confirm this effect in a neuronal context, ROS levels were quantified in RA-differentiated SH-SY5Y cells by flow cytometry (Figure 3D). Forward- and side-scatter gating was used to define the main cell population and exclude debris (Figure S6B). Tunicamycin-treated cells displayed increased CM-H₂DCFDA fluorescence, whereas co-treatment shifted the fluorescence distribution leftward and reduced the median fluorescence intensity (MFI), consistent with lower ROS levels (Figures 3E–3F and S6A). This effect was reproduced in an independent biological replicate, with a consistent reduction observed at 1.6 μM (Figures S6C–S6D). Together, these data indicate that SR-0813 limits oxidative stress associated with ER dysfunction.

**Figure 3.**
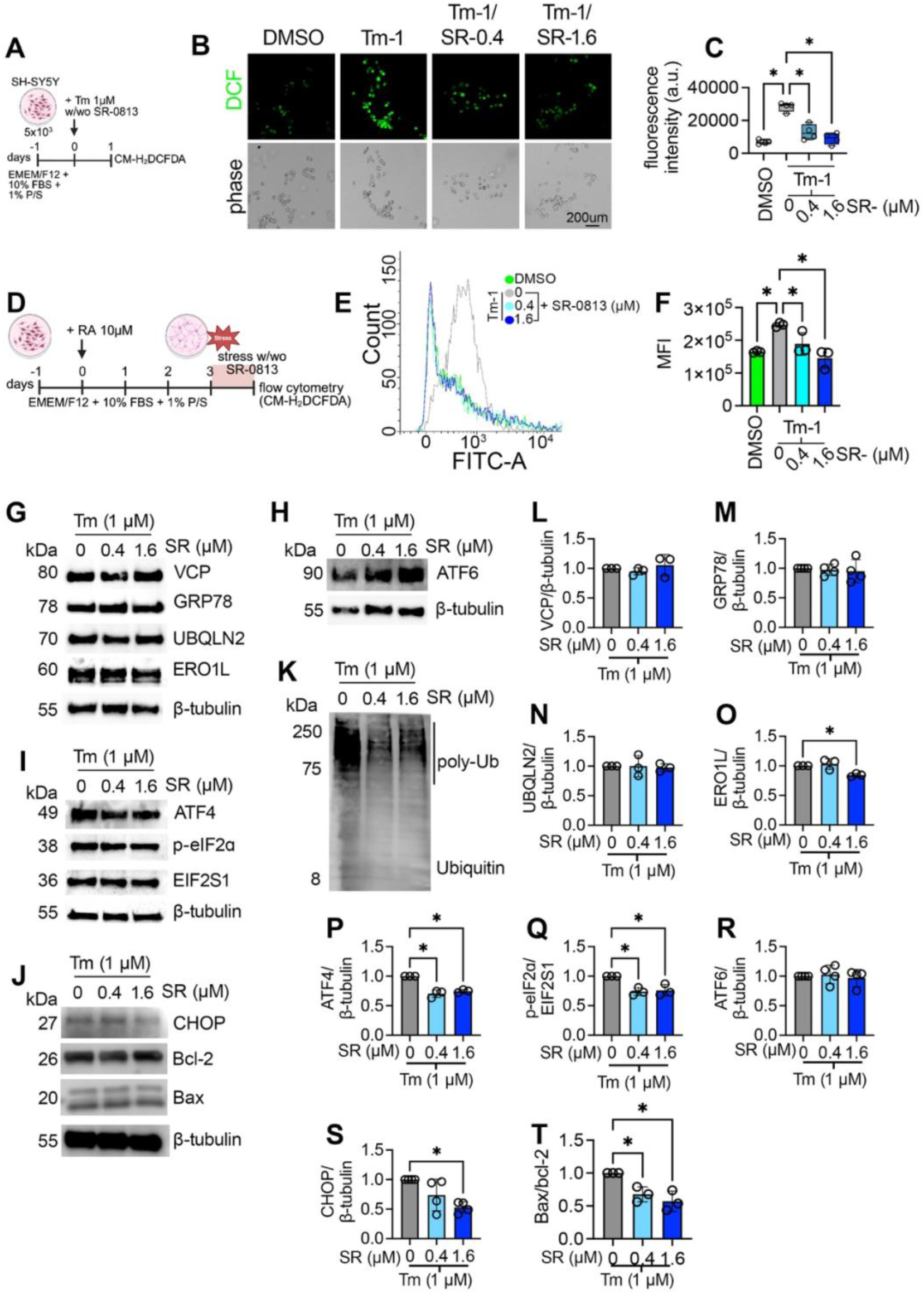
SR-0813 reduces tunicamycin-induced ROS and dampens ER stress/ISR and apoptotic signaling. (A–C) Confocal ROS imaging in undifferentiated SH-SY5Y cells. Cells were exposed to tunicamycin (Tm; 1 µM) and co-treated with vehicle (DMSO) or SR-0813 (0.4 or 1.6 µM). Intracellular ROS was measured using CM-H₂DCFDA. (B) Representative CM-H₂DCFDA fluorescence images (top) and corresponding phase-contrast images (bottom). (C) Quantification of CM-H₂DCFDA fluorescence intensity (a.u.) shown as box-and-whisker plots with all data points overlaid; whiskers indicate the full range (min to max). (D–F) Flow-cytometric ROS measurements in differentiated SH-SY5Y cells. (D) Experimental design: differentiated cells were treated with Tm and co-treated with SR-0813 (0, 0.4, or 1.6 µM) or left unstressed (DMSO), then stained with CM-H₂DCFDA and analyzed by flow cytometry. (E) Representative histograms (count versus FITC-A). (F) Quantification of median fluorescence intensity (MFI). Points represent biological replicates from 3 independent experiments (n = 3); bars show mean ± SD. Statistical analysis was performed by one-way ANOVA with Brown–Forsythe correction; p < 0.05 was considered significant. (G–K) Immunoblot analysis of ER stress/ integrated stress response (ISR), proteostasis, and apoptosis markers in SH-SY5Y cells treated as in (D). β-tubulin was used as a loading control. (L–T) Densitometric quantification of immunoblots in (G–K) from 3 independent experiments (n = 3). Values were normalized to β-tubulin, except p-eIF2α normalized to total eIF2α (EIF2S1); Bax/Bcl-2 is presented as a ratio. Points represent biological replicates; bars show mean ± SD. SR-indicates SR-0813. Statistical significance was assessed by ordinary one-way ANOVA with Sidak’s multiple-comparisons test; p < 0.05 was considered significant.

We next investigated whether this reduction in oxidative stress was accompanied by modulation of ER stress signaling pathways. Immunoblot analysis revealed that expression of the ER chaperone GRP78/BiP was not altered by co-treatment (Figures 3G and 3M). In contrast, levels of ERO1L, an oxidoreductase that contributes to oxidative protein folding within the ER, were reduced at the higher concentration (1.6 μM) (Figures 3G and 3O).

To further dissect UPR signaling, we examined distinct branches of the response. ATF6 levels were not affected by co-treatment (Figure 3H and 3R), indicating that this arm of the UPR remains largely unchanged. By contrast, markers of the PERK–ISR axis were selectively attenuated. Co-treatment reduced ATF4 levels and decreased the ratio of phosphorylated eIF2α to total eIF2α, consistent with suppression of ISR signaling (Figures 3I and 3P–3Q). This was accompanied by a reduction in CHOP levels at 1.6 μM (Figures 3J and 3S), indicating dampening of the maladaptive, pro-apoptotic branch of the UPR. In line with this, the Bax/Bcl-2 ratio was decreased (Figures 3J and 3T), supporting reduced apoptotic commitment, consistent with the reduced apoptosis observed by Annexin V/propidium iodide staining (Figures 2F–2M).

Given the link between ER stress and proteostasis impairment, we next examined the accumulation of ubiquitinated proteins. Co-treatment reduced the levels of polyubiquitinated (poly-Ub) species under Tm stress (Figure 3K), suggesting improved protein quality control capacity.

Finally, to determine whether these effects reflect a broader modulation of proteostasis pathways, we assessed additional proteins involved in ER homeostasis and protein quality control. The abundance of VCP and UBQLN2 was not altered by co-treatment under tunicamycin stress (Figures 3G and 3L–3N), indicating that the observed effects occur without widespread changes in these proteostasis-associated factors.

To determine whether these effects extend beyond ER stress conditions, we performed a parallel analysis in cells exposed to proteasomal inhibition (MG132), where the protective effects were more modest (Figure 2). Given that MG132 primarily perturbs proteostasis rather than directly engaging canonical UPR signaling, we focused this analysis on protein quality control and apoptotic markers.

Under these conditions, co-treatment did not alter VCP or UBQLN2 levels (Figure S7A–S7C), consistent with the findings observed under Tm stress. Accumulation of poly-Ub proteins was only mildly reduced and reached significance at the higher concentration (1.6 μM) (Figure S7A), indicating a limited impact on proteostasis under these conditions.

Despite this, markers of apoptotic signaling were consistently attenuated, with reduced CHOP levels and a lower Bax/Bcl-2 ratio observed upon co-treatment (Figure S7D–S7E). Thus, while the effects under proteasomal stress are less pronounced than those observed under ER stress, the compound retains the ability to modulate downstream stress signaling and apoptotic commitment without altering core proteostasis regulators.

Together, these results indicate that SR-0813 reduces ER stress-associated oxidative stress and PERK-ISR signaling, limiting apoptosis and improving proteostasis. Its effects under proteostasis impairment are more modest but consistent, without broadly affecting core proteostasis regulators.

### ENL/AF9 modulation exerts context-dependent effects across neurodegenerative models with selective impact on UBQLN2-associated pathology

Building on the observation that SR-0813 enhances neuronal resilience under ER stress, we next sought to determine whether these effects reflect modulation of ENL/AF9 itself and to what extent this pathway contributes across distinct neurodegenerative contexts. To address this, we complemented our chemical approach with a genetic strategy in *Drosophila*, enabling direct interrogation of ENL/AF9 function independent of compound-specific effects and across multiple *in vivo* disease models.

To this end, we utilized a panel of transgenic *Drosophila* models expressing human neurodegeneration-associated proteins in the eye under the control of the GMR-GAL4 driver, which directs expression to differentiating photoreceptor neurons. In this system, expression of disease-associated proteins produces a readily quantifiable rough eye phenotype, providing a sensitive *in vivo* readout of neuronal toxicity and genetic modification. The panel encompassed models of major neurodegenerative diseases, including amyotrophic lateral sclerosis (ALS), Alzheimer’s disease, Huntington’s disease, Parkinson’s disease, and spinocerebellar ataxia. Notably, several of these disease-associated proteins have been linked to ER stress, although the underlying mechanisms are heterogeneous and remain incompletely defined ^20^.

Flies expressing these human transgenes were analyzed in the presence or absence of *dENL/AF9* knockdown, where control animals carried a non-targeting RNAi construct to ensure a matched genetic background (GMR-GAL4 driving two UAS transgenes). Eye degeneration was scored relative to these controls (Figures 4A-E).

**Figure 4.**
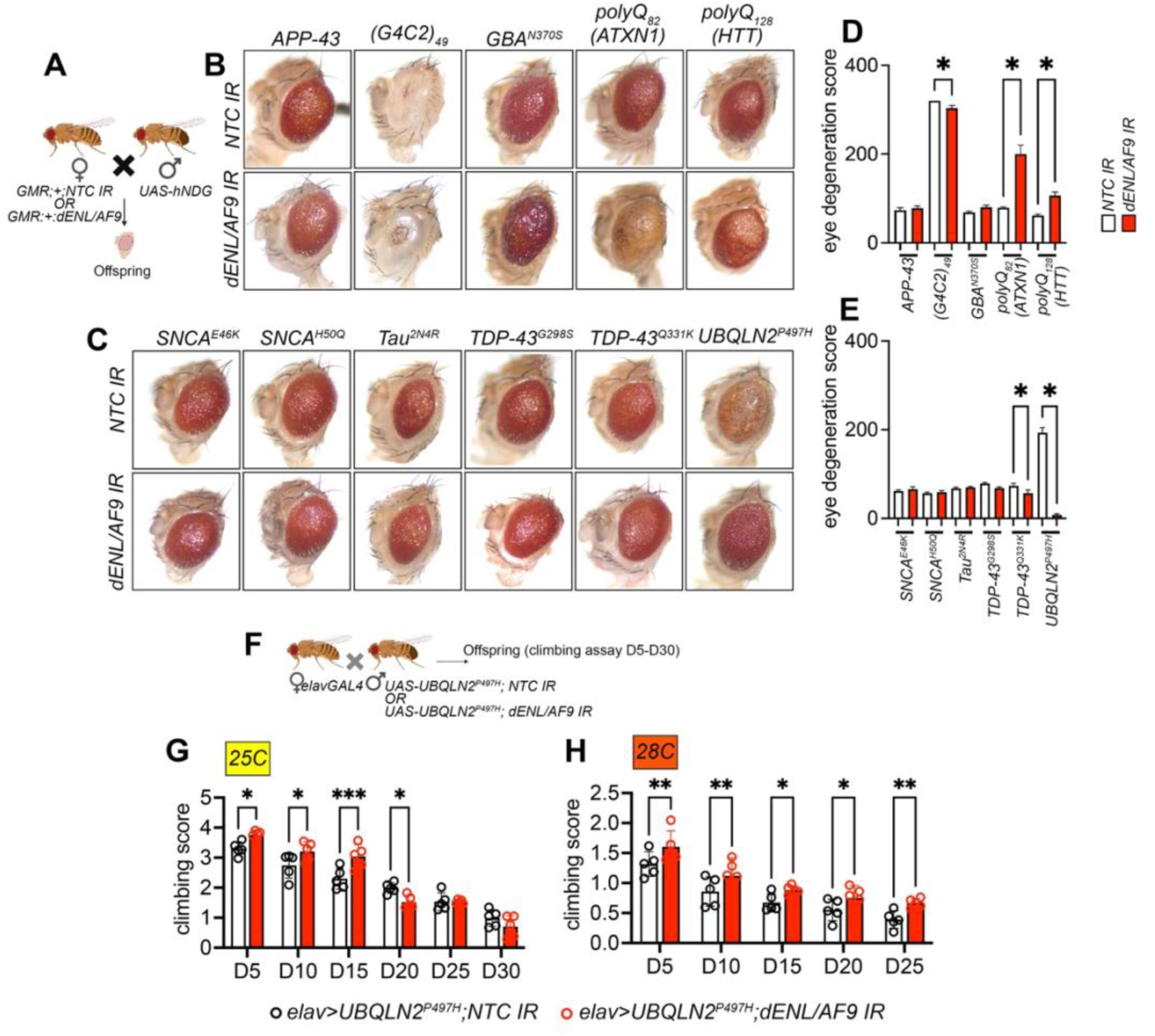
dENL/AF9 knockdown is a model-dependent modifier of neurodegenerative phenotypes. (A) Schematic of the genetic modifier screen testing knockdown (KD) of the *Drosophila* ENL/AF9 homolog (dENL/AF9) across fly models of neurodegeneration. (B–C) Representative adult eye images from flies expressing human disease-associated proteins in the eye under the GMR-GAL4 driver, with or without dENL/AF9 KD. Models included Alzheimer’s disease/amyloid pathology (APP-43), tauopathy (Tau2N4R), Parkinson’s disease (GBA^N370S^, SNCA^E46K^, SNCA^H50Q^), polyglutamine disorders (ATXN1 polyQ82; HTT polyQ128), and ALS/FTD (G4C2 repeats, TDP-43^Q331K^, TDP-43^G298S^, and UBQLN2^P497H^). Eye degeneration (“rough eye”) served as the primary screening readout. (D–E) Quantification of rough-eye degeneration scores for each disease model expressed under GMR-GAL4 and co-expressed with either dENL/AF9 RNAi or a non-targeting control (NTC) inverted repeat (IR). Bars show mean ± SD. (F) Experimental design for assessing dENL/AF9 KD in a disease-relevant tissue. Pan-neuronal expression was driven by elav-GAL4, and flies were reared at 25°C or 28°C to modulate GAL4/UAS activity (KD efficiency at 25 and 28°C is shown in Figure S7). (G) Negative geotaxis (climbing) performance for the indicated genotypes (elav-GAL4) plotted as climbing score based on distance climbed at 5, 10, 15, 20, 25, and 30 days post-eclosion. Bars show mean ± SD; a total of 100 flies were analyzed per genotype and time point. For (D–E) and (G), statistical significance was determined by one-way ANOVA with Dunnett’s multiple-comparisons test; p < 0.05 was considered significant. (hNDG: human neurodegenerative gene; NTC: non-targeting gene)

Genetic reduction of *dENL/AF9* produced context-dependent effects across neurodegenerative models. A robust suppression of retinal degeneration was observed in flies expressing *UBQLN2^P^*^497^*^H^*, a mutation associated with familial ALS (Figures 4C-E). More modest improvement was detected in TDP-43^Q331K^ and (G4C2)_49_ repeat models, whereas toxicity was exacerbated in polyglutamine disease models (polyQ), including ATXN1-polyQ_82_ and HTT-polyQ_128_ (Figures 4A-E).

The pronounced rescue of UBQLN2-associated toxicity is consistent with our cellular findings, where ENL/AF9 modulation attenuated ER stress and PERK-ISR signaling, pathways that are strongly implicated in UBQLN2-linked disease. In contrast, as ployQ toxicity is driven by a persistent burden of aggregation-prone proteins that requires sustained chaperone, autophagy, and proteasome activity, these findings suggest that ENL/AF9 inhibition may be beneficial when pathology is dominated by stress-pathway signaling and proteotoxic load, but detrimental when long-term aggregation-buffering capacity is the limiting factor.

Given the pronounced rescue observed in the UBQLN2^P497H^ model, we next sought to evaluate this effect in a more disease-relevant neuronal context. To this end, we generated a recombinant fly line carrying both elav-GAL4 and *UAS-dENL/AF9 RNAi*, enabling pan-neuronal knockdown of *dENL/AF9* (Figure S8A). Knockdown efficiency was modest under standard conditions and was slightly enhanced at 28°C, consistent with the temperature sensitivity of the UAS–GAL4 system ^21^, although remaining overall partial (Figures S8B-D).

We then assessed locomotor function in flies expressing UBQLN2^P497H^ in neurons in the presence or absence of *dENL/AF9* knockdown (Figures 4F-H). At 25°C, *dENL/AF9* reduction improved climbing performance at early stages of disease progression, with limited benefit at later time points. Increasing the temperature to 28°C, which enhances GAL4-driven knockdown, resulted in a modest but more sustained improvement across disease progression. Notably, UBQLN2^P497H^ flies exhibited a more severe phenotype at 28°C, which may constrain the overall magnitude of the observed rescue.

Together, these findings support a context-dependent role for ENL/AF9 in regulating neuronal stress responses and vulnerability, with selective protective effects in UBQLN2-associated pathology.

### SR-0813 ameliorates neuronal dysfunction and stress signaling in a *Drosophila* UBQLN2-ALS model

Having established that genetic reduction of *dENL/AF9* confers protection in UBQLN2-associated pathology, we next tested whether pharmacological inhibition could recapitulate these effects *in vivo*. SR-0813 was administered throughout development to *Drosophila* expressing UBQLN2^P497H^, and neuronal function and pathology were assessed.

Pan-neuronal expression of UBQLN2^P497H^ in *Drosophila* larvae leads to measurable locomotor (crawling) impairments ^20^. We therefore asked whether this phenotype could be modified by SR-0813 treatment. Larvae were therefore exposed to SR-0813 (1 or 10 μM) from early developmental stages, and locomotor behavior was evaluated at the third instar stage (Figure 5A). Treatment resulted in a robust improvement in crawling performance, with increased path length, distance traveled and average speed compared to untreated UBQLN2^P497H^ larvae (Figures 5B–5E).

**Figure 5.**
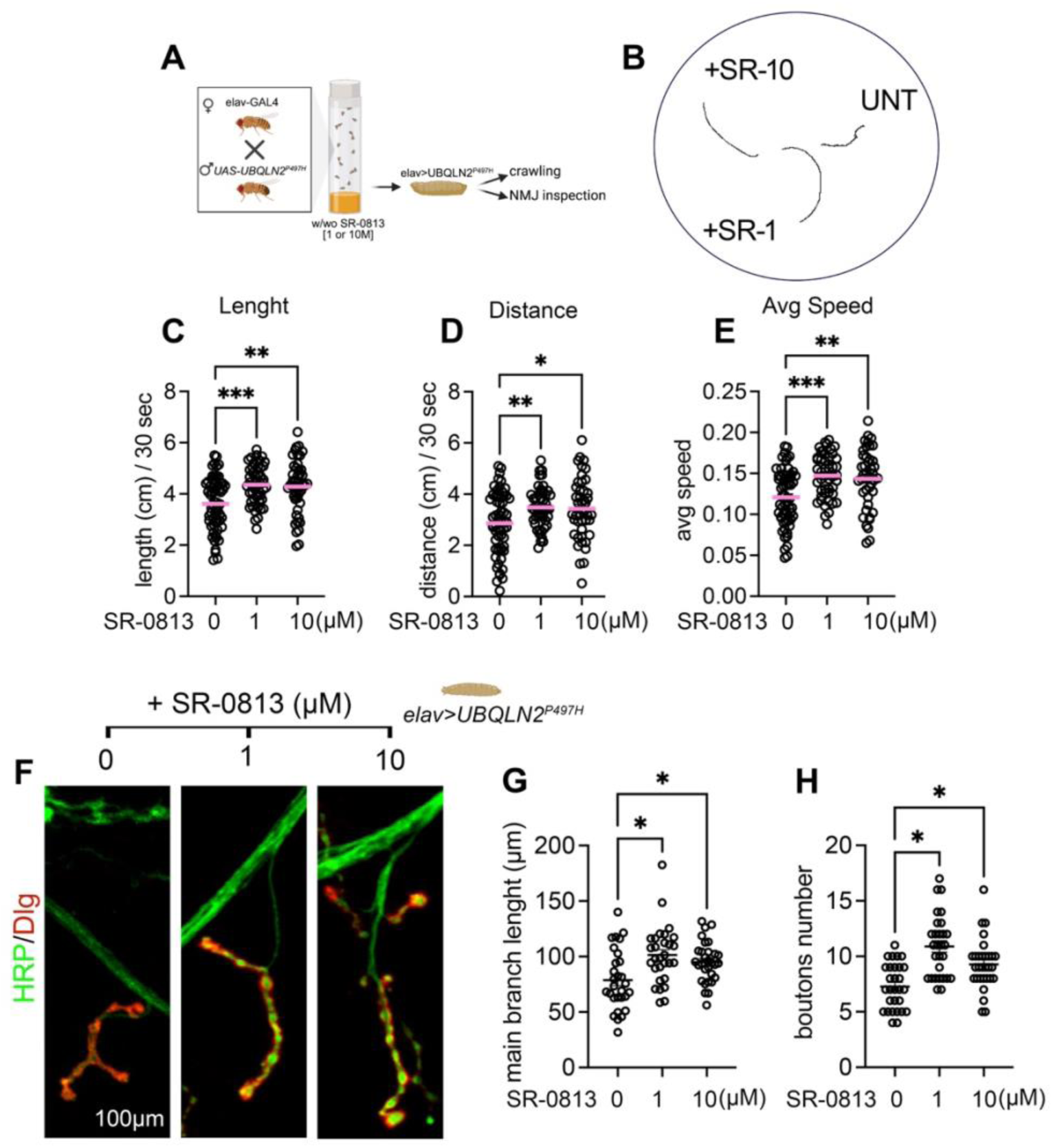
SR-0813 improves locomotor function and neuromuscular junction morphology in a *Drosophila* UBQLN2^P497H^ ALS model. (A) Experimental design. UBQLN2^P497H^ was expressed pan-neuronally using elav-GAL4. Parental flies were maintained on standard food supplemented with SR-0813 (1 µM; SR-1 or 10 µM; SR-10) or left untreated (UNT). Progeny were assessed at the third-instar larval stage (L3) by crawling assays, and adults were assayed for climbing performance every 5 days from day 5 to day 30 post - eclosion (D5-D30). (B–E) Larval crawling behavior. (B) Representative 30-s locomotor tracks from L3 foraging larvae (UNT, SR-1, SR-10). Quantification of (C) track length, (D) distance traveled, and (E) average speed over 30 s. Each point represents one larva (UNT n = 60; SR-1 n = 53; SR-10 n = 57). Statistical analysis: ordinary one-way ANOVA with Dunnett’s multiple-comparisons test (each SR-0813 condition versus UNT); p < 0.05 was considered significant. (F–H) Neuromuscular junction (NMJ) morphology. (F) Representative images of type Ib NMJs on muscle 4 (abdominal segment A4) from L3 elav>UBQLN2^P497H^ larvae reared on UNT, SR-1, or SR-10 food. Presynaptic terminals were labeled with anti-HRP (green) and postsynaptic densities with anti-Discs large (Dlg; red). Quantification of (G) NMJ branch length and (H) bouton number. Each point represents one larva (n = 30 larvae per condition). Statistical analysis: ordinary one-way ANOVA with Dunnett’s multiple-comparisons test (each SR-0813 condition versus UNT); p < 0.05 was considered significant.

To determine whether this functional improvement was accompanied by structural preservation, we analyzed neuromuscular junction (NMJ) morphology, which is known to be compromised upon pan-neuronal UBQLN2^P497H^ expression in flies ^22,23^. SR-0813 treatment increased NMJ branch length and bouton number at muscle 4 in abdominal segment A4, indicating partial restoration of synaptic architecture (Figures 5F–5H).

We next examined whether these effects were associated with modulation of disease-relevant molecular pathways *in vivo*. Oxidative stress was assessed in adult brains of UBQLN2^P497H^ at 15 days of age (Figure 6A), a stage characterized by progressive neurodegeneration and locomotor decline (Figures 4G–4H and S9). Dihydroethidium (DHE) staining revealed that treatment reduced ROS levels in UBQLN2^P497H^ fly brains, as evidenced by decreased fluorescence intensity and ROS index, particularly at 10 μM (Figures 6A–6C). This effect is consistent with the attenuation of oxidative stress observed in neuronal cell models (Figures 3A–3F). Notably, UBQLN2^P497H^ protein levels remained unchanged upon treatment, indicating that the observed rescue is not attributable to altered transgene expression or turnover (Figure 6D-E).

**Figure 6.**
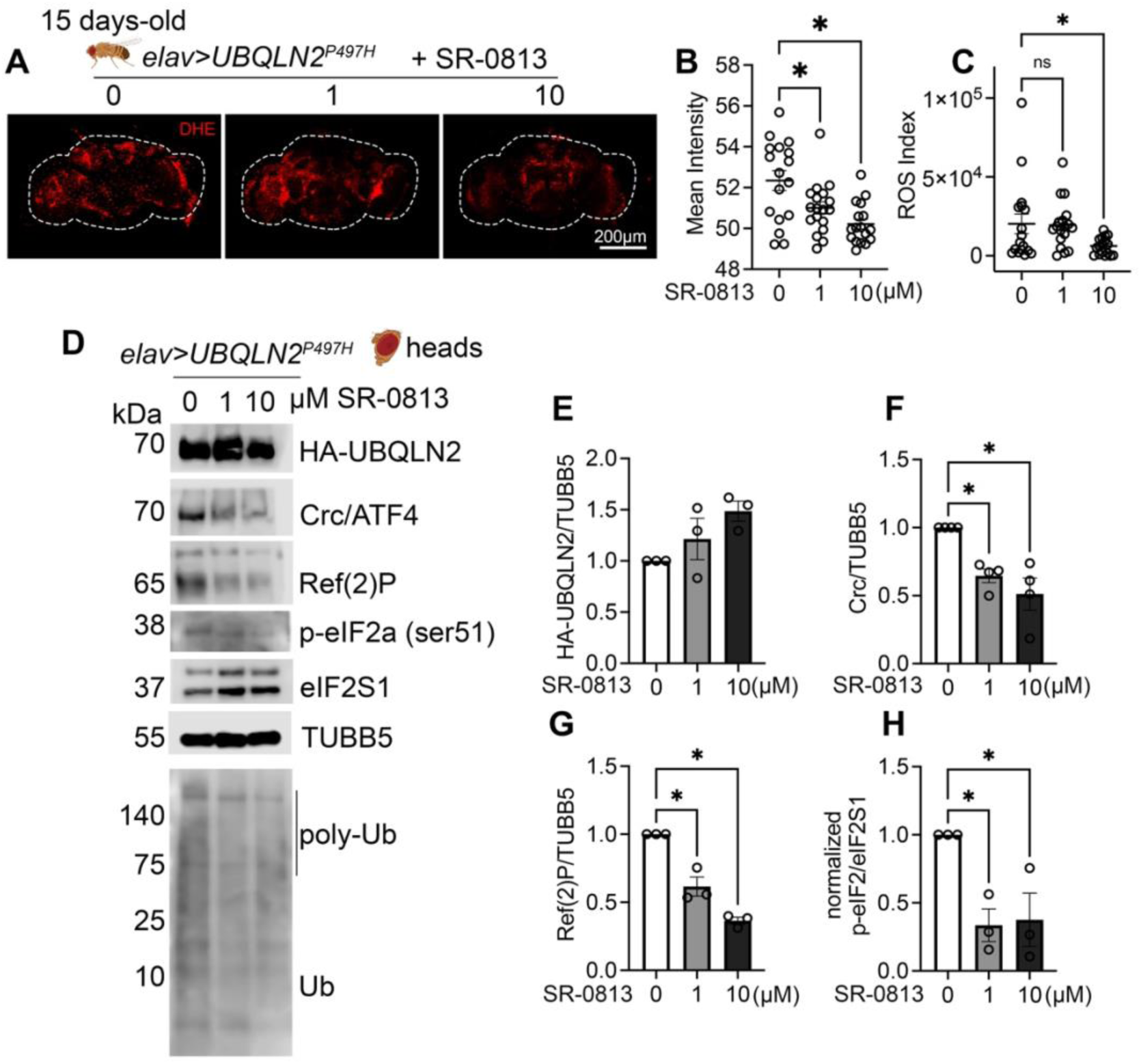
SR-0813 reduces oxidative stress and dampens ISR/proteostasis stress markers in UBQLN2^P497H^ flies. (A) Representative confocal images of adult brains stained with dihydroethidium (DHE) to assess ROS in UBQLN2^P497H^ flies left untreated (UNT) or treated with SR-0813 (1 µM, SR-1; 10 µM, SR-10). Flies were reared on standard food and, after eclosion, maintained on SR-0813–supplemented food (or UNT) for 15 days prior to imaging. (B–C) Quantification of brain ROS. (B) Mean DHE fluorescence intensity and (C) ROS index, defined as *total intensity × (N objects / mean N objects in the control group)*, quantified using CellSense (Olympus FLUOVIEW FV300). Each point represents one brain (n = 17). Statistical analysis: ordinary one-way ANOVA with Sidak’s multiple-comparisons test; p < 0.05 was considered significant. (D) Immunoblot analysis of head lysates from UBQLN2^P497H^ flies treated as in (A). HA-UBQLN2 was measured to confirm that phenotypic rescue was not attributable to reduced UBQLN2 expression. Markers of the integrated stress response (ISR) included *Drosophila* ATF4 (Cryptocephal, Crc) and phospho-eIF2α (p-eIF2α) with total eIF2α (eIF2S1). Ref(2)P and ubiquitin (polyubiquitinated species) were assessed as proteostasis/autophagy-associated readouts. β-tubulin (TUBB) served as a loading control. (E–H) Densitometric quantification of immunoblots in (D) from 3 independent experiments (n = 3). (E) HA-UBQLN2, (F) Crc, (G) Ref(2)P, and (H) p-eIF2α normalized to total eIF2α (eIF2S1). Signals were normalized to β-tubulin (TUBB) unless otherwise indicated. Points represent biological replicates; bars show mean ± SD. Statistical analysis: ordinary one-way ANOVA with Dunnett’s multiple-comparisons test (each SR-0813 condition versus UNT); p < 0.05 was considered significant.

We then assessed ER stress and proteostasis-related pathways. Treatment reduced levels of the ISR transcription factor ATF4 (cryptocephalus, Crc in flies) and phosphorylation of eIF2α (Ser51), indicating attenuation of PERK–ISR signaling *in vivo*, consistent with our findings in human neurons (Figure 6D-H). In parallel, levels of the autophagy adaptor Ref(2)P and of polyubiquitinated protein species were reduced in a dose-dependent manner, consistent with decreased accumulation of undegraded autophagic cargo and a lower burden of misfolded proteins, suggestive of an overall enhancement of autophagic/proteostatic clearance (Figures 6D, 6G).

Together with the absence of detectable changes in the proteostasis factors assessed *in vitro*, these findings suggest that ENL/AF9 inhibition reduces proteotoxic burden, coincident with dampened ISR signaling.

Collectively, these findings demonstrate that pharmacological inhibition of ENL/AF9 recapitulates genetic protection *in vivo*, improving neuronal function and attenuating oxidative and ER stress signaling in a UBQLN2-associated ALS model.

### SR-0813 differentially modulates locomotor decline across *Drosophila* ALS models

Building on the earlier eye-based genetic modifier analysis, which suggested beneficial interactions in multiple ALS-linked contexts, we next asked whether pharmacological inhibition could confer protection across diverse ALS models. To address this, we evaluated the effects of SR-0813 in a panel of *Drosophila* lines expressing ALS-associated proteins.

We first performed a preliminary characterization of locomotor dysfunction to establish a platform spanning a range of disease severities and genetic etiologies (Figure S8). Pan-neuronal expression of (G4C2)_49_, TDP-43^Q331K^, TDP-43^G298S^, UBQLN2^P497H^, and SOD1 mutants using elav-GAL4 revealed a spectrum of age-dependent impairments. TDP-43^Q331K^ induced more pronounced deficits than TDP-43^G298S^, whereas SOD1^G94A^ and SOD1^A4V^ exhibited comparatively milder, progressive decline. Based on these profiles and their translational relevance, we selected (G4C2)_49_, TDP-43^Q331K^, SOD1^G94A^, and UBQLN2^P497H^ for further analysis.

SR-0813 was administered from early developmental stages, and locomotor performance was assessed longitudinally from day 5 to day 30 post-eclosion (Figure 7). Consistent with earlier findings, SR-0813 maintained a robust protective effect in UBQLN2^P497H^ flies, improving climbing performance across disease progression, including at later stages (Figure 7A). In SOD1^G94A^ flies, treatment conferred a significant benefit at later time points, particularly beyond day 15, when locomotor impairment becomes more pronounced (Figure 7B and Figure S8). A modest but significant improvement was also observed in TDP-43^Q331K^ flies, predominantly at later stages, although treatment did not prevent premature lethality beyond 20 days (Figure 7C).

**Figure 7.**
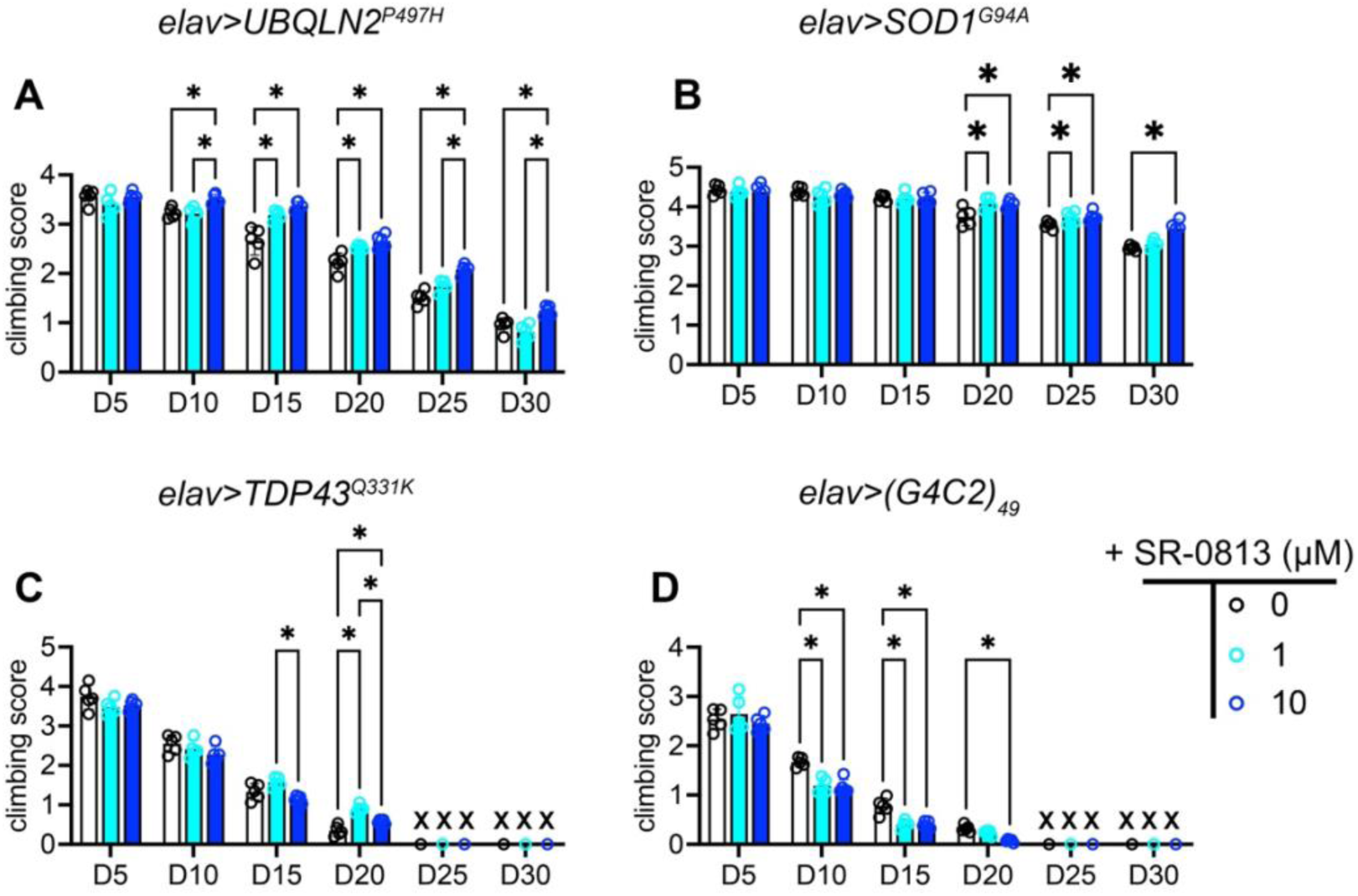
SR-0813 differentially modulates adult locomotor decline across *Drosophila* ALS models. Adult locomotor function was assessed longitudinally across multiple Drosophila ALS models to determine whether SR-0813–mediated effects were sustained during adulthood and across disease progression. Human ALS-associated genes were expressed pan-neuronally using elav-GAL4, and flies were maintained on standard food supplemented with SR-0813 (1 µM or 10 µM) or left untreated (UNT). Food was replaced every 5 days, and negative geotaxis (climbing) performance was measured every 5 days from 5 to 30 days post-eclosion (D5–D30). “x” marks indicate time points at which flies died. (A–D) Climbing performance of (A) UBQLN2^P497H^, (B) SOD1^G94A^, (C) TDP-43^Q331K^, and (D) (G4C2)_49_ flies under UNT, SR-0813 (1 µM), or SR-0813 (10 µM) conditions. Data are shown as bar plots with individual values from 5 independent replicates (biological replicate vials) at each time point; a total of 100 flies were tested per genotype/condition. Statistical analysis was performed by ordinary one-way ANOVA with Dunnett’s multiple-comparisons test (each SR-0813 condition versus UNT); p < 0.05 was considered significant. Climbing performance relative to driver/background controls is shown in Figure S8. Black “x” symbols indicate time points at which the corresponding genotype had died (no climbing measurement possible).

In contrast, SR-0813 exacerbated locomotor decline in flies expressing (G4C2)_49_ repeats (Figure 7D), indicating that ENL/AF9 inhibition exerts context-dependent effects across ALS models. Notably, this effect emerged with age, as treatment did not significantly alter locomotor performance in young flies but worsened decline as the flies get old.

Overall, this pattern is consistent with SR-0813 reducing proteotoxic stress burden and ISR signaling, conferring benefit in UBQLN2^P497H^ and SOD1^G94A^ models, but proving less effective or detrimental when pathology is dominated by sustained aggregation burden requiring long-term proteostasis support.

## DISCUSSION

ENL/AF9 YEATS-domain proteins are best known as chromatin readers that promote transcriptional elongation, yet their roles in neuronal stress adaptation have been unclear. Building on our prior finding that dENL/AF9 depletion extends *Drosophila* lifespan and improves stress tolerance, we used these phenotypes as benchmarks to test whether the selective YEATS inhibitor SR-0813 elicits similar organismal outcomes, consistent with functional engagement of the ENL/AF9 pathway in flies. *In vivo*, SR-0813 phenocopied key aspects of *dENL/AF9* knockdown: late-onset treatment extended lifespan, enhanced resistance to oxidative challenge, and ameliorated locomotor decline and NMJs defects in a UBQLN2-linked ALS model. These convergent genetic and pharmacologic data support a conserved role for ENL/AF9 in shaping stress resilience at the organismal level and motivated deeper mechanistic studies in neuronal models.

SR-0813 showed no measurable radical-scavenging activity in cell-free assays, suggesting its ROS-lowering effects in cells and *in vivo* may arise indirectly from YEATS inhibition reshaping stress signaling. In cellular models, SR-0813 dampened the PERK/eIF2α/ATF4/CHOP axis and reduced apoptotic outputs without broadly preventing UPR induction or altering the proteostasis factors examined. These findings indicate that SR-0813 primarily tunes stress-response signaling rather than globally increasing proteostasis capacity, consistent with its ability to mitigate degeneration in a UBQLN2-linked ALS model in which toxicity has been associated with amplified ISR engagement ^24,25^.

The parallel phenotypes produced by YEATS inhibition and genetic knockdown further suggest that ENL/AF9 activity can be modulated through distinct modalities, acutely by disrupting reader function or chronically by reducing expression, highlighting opportunities for future studies using orthogonal YEATS ligands or ENL-directed degradation strategies ^26–28^ to more comprehensively define ENL/AF9 function in neuronal contexts.

A striking feature of ENL/AF9 inhibition was its context dependence. SR-0813 was protective in paradigms driven by acute ER stress or maladaptive ISR activation, but showed reduced efficacy and in some cases detrimental effects in models associated with sustained mitochondrial dysfunction or aggregation stress, including rotenone exposure and polyglutamine- or repeat-expansion-driven toxicity. These chronic conditions impose a persistent demand for adaptation, requiring continuous engagement of mitochondrial quality control, autophagy, and other proteostasis-support pathways ^29–31^.

These divergent outcomes are consistent with a trade-off in which ENL/AF9 inhibition is beneficial when it limits excessive ISR output, but becomes unfavorable when long-term survival depends on maintaining transcriptional capacity for sustained adaptation. Such a balance may be particularly relevant for aggregation-prone proteins, where toxicity is strongly shaped by chaperone availability and clearance pathways, and for mitochondrial stress, where ongoing compensatory programs are required to preserve bioenergetic homeostasis ^32–34^.

This framework aligns with the molecular function of ENL/AF9 in transcriptional elongation complexes, where YEATS-domain-dependent recognition of histone acyl-lysine marks supports RNA polymerase II pause release and amplifies stimulus-responsive transcription ^35,36^. In this view, the YEATS domain couples chromatin engagement to elongation control: acyl-lysine recognition promotes locus-specific recruitment and/or stabilization of ENL/AF9, enabling ENL/AF9-containing elongation complexes to increase transcriptional output. Because histone acylation is metabolically sensitive ^37,38^, ENL/AF9 may thereby link stress-and metabolism-driven chromatin states to the magnitude and persistence of stress-induced transcriptional programs. We therefore propose that ENL/AF9 inhibition modulates the amplitude and persistence of stress-induced transcriptional responses, with outcomes determined by the nature and duration of the insult.

Notably, other YEATS-domain readers appear to support distinct neuronal transcriptional functions; for example, YEATS2, the reader subunit of the ATAC histone acetyltransferase complex, has been implicated in ribosomal and metabolic gene expression ^39,40^, and its depletion causes neuronal stress and behavioral defects in *Drosophila* ^17,41^, consistent with a role in maintaining basal biosynthetic capacity. In this light, our data support a model in which YEATS readers are specialized in neurons, with YEATS2 safeguarding core biosynthetic programs and ENL/AF9 tuning stress-responsive gene amplitude.

Overall, these findings extend ENL/AF9 biology beyond its established role in transcriptional control and its current therapeutic focus in cancer, identifying YEATS-domain inhibition as a means to modulate neuronal stress adaptation. The divergent effects observed across models indicate that ENL/AF9 targeting does not confer uniform cytoprotection in neurons; rather, its impact depends on the dominant stress mechanism within a given context-beneficial when pathology is driven by excessive ISR and apoptotic signaling, but less effective or even detrimental when long-term survival relies on sustained proteostasis and metabolic adaptation. Defining the stress conditions and disease settings in which this balance can be favorably shifted will be critical for evaluating the therapeutic potential of ENL/AF9 inhibition in neurodegenerative disorders.

### Limitations of the study

This study has several limitations that point to important next steps. First, although SR-0813 is a well-characterized, chromatin-competitive ENL/AF9 YEATS inhibitor, we did not directly quantify its target engagement or pharmacokinetics in Drosophila or mammalian systems, and more detailed exposure and occupancy measurements will be needed to refine dosing windows. Second, our mechanistic work was performed in RA-differentiated SH-SY5Y cells and Drosophila neurons; extending these findings to human iPSC-derived motor neurons and mammalian ALS models will be important to test the generality and translational relevance of the context-dependent effects we observe. Finally, we focused on defined ISR, proteostasis, and apoptotic markers and did not undertake genome-wide chromatin or transcriptomic profiling, so the full spectrum of ENL/AF9-regulated stress-response programs remains to be mapped.

## RESOURCE AVAILABILITY

### Lead contact

Requests for further information and resources should be directed to and will be fulfilled by the lead contact, Luca Lo Piccolo

### Materials availability

All unique/stable reagents generated in this study are available from the lead contact with a completed materials transfer agreement.

### Data and code availability

All data supporting the findings of this study are available from the corresponding author upon reasonable request. No custom code was used for data acquisition or analysis beyond standard software and publicly available packages described in the Methods.

## ACKNOWLEDGMENTS

This work was supported by the Fundamental Fund (FF) from the Science, Research and Innovation Promotion Fund (TSRI), Chiang Mai University, FY2568 under Contract No. 207569. This study was partially supported by the FF, FY2568 Contract No. 207573, the Faculty of Medicine, Chiang Mai University grant no FACMED 139/2567, Health Systems Research Institute (HSRI) under grants 66-124 and 68-060, the Mid-Career Research Grant (N42A670768) of the National Research Council of Thailand and PM and other Pollutants Related NCDs from Field to Cell to Bedside (FCB) funding, Chiang Mai University under the grants PM14/2566. We thank Mr. Papon Hitmool, Mr. Sakorn Phongchankhiao, and Mr. Tawan Munwanna (Department of Pharmacology, Faculty of Medicine, Chiang Mai University) for their technical and administrative assistance. We thank Dr. Yoshida Hideki (Kyoto Institute of Technology) and the Bloomington Stock Center for providing fly strains, the Science and Educational Company Limited (SCIED) for training in confocal scanning electron microscopy.

## AUTHOR CONTRIBUTIONS

Conceptualization: L.LP. and S.J.; Data curation: L.LP. and S.J.; Formal Analysis: L.LP., C.P., Ra.Y., Ru.Y. and S.J.; Funding acquisition; L.LP.; Investigation: L.LP and S.J.; Methodology: L.LP., Ra.Y., Ru.Y and S.J; Project administration: L.LP.; Resources; L.LP and S.J; Software: L.LP.; Supervision; L.LP; Validation: L.LP. and S.J.; Visualization: L.LP; Writing – original draft: L.LP.; Writing – review & editing: All authors have reviewed and approved the final version of this manuscript.

## SUPPLEMENTAL INFORMATION

**Document S1. Figures S1–S13, Tables S1-S4**

## Materials and Methods

### *Drosophila* stocks and husbandry

*Drosophila melanogaster* stocks were maintained on a standard cornmeal-yeast-glucose medium at 22°C with 60% humidity under a 12:12 light/dark cycle unless otherwise stated. All *Drosophila* stocks and full genotypes used in this study, including GAL4 drivers, UAS-RNAi lines, and neurodegenerative disease model transgenes, were obtained from the Bloomington Drosophila Stock Center (BDSC). GMR-Gal4 (X) and *GMR-Gal4;+;UAS-GFP* IR are a gift of Prof. Yoshida (Kyoto Institute of Technology). A full list of fly lines used in this study is provided in Table S1. The Gal4/UAS binary system was used for targeted, tissue-specific knockdown or overexpression ^42^. Crosses were performed at 25°C unless indicated; for experiments requiring increased GAL4 activity, flies were maintained at 28°C as described. Adult flies were collected within 24 h of eclosion and aged under standard density conditions. Pan-neuronal knockdown or overexpression was achieved using the elav-GAL4 drivers at 25°C. The glass multiple reporter (GMR)-GAL4 promoter was used for targeting transgene’s expression to developing eyes at 28°C. To minimize genetic background effects, all experimental flies were backcrossed six times with the *w* strain before use.

### Compounds and preparation

All compounds were initially prepared as concentrated stock solutions in 100% DMSO and diluted to working concentrations immediately before use, except where noted. Retinoic acid and tunicamycin stocks were prepared at 100 mM and 5 mg/mL, respectively. The selective ENL/AF9 YEATS-domain inhibitor SR-0813 was prepared as 20 mM stock solutions in DMSO. For cell-based experiments, compounds were applied at final concentrations and durations indicated in the figure legends. For fly experiments, SR-0813 was administered by supplementation into food at the indicated concentrations. Unless otherwise specified, drug-containing food was replaced every 5 days. Details of all compounds and their CAS registry numbers are provided in Supplementary Table S2.

### Measurement of radical scavenging activity

Radical scavenging activity of compounds was assessed using complementary DPPH and ABTS assays, as previously described ^43^. Briefly the DPPH assay measured neutralization of lipophilic free radicals in methanolic solution, while the ABTS assay evaluated both hydrophilic and lipophilic antioxidant activity in aqueous conditions. For DPPH, a 0.2 mM solution in methanol was incubated with test compounds in 96-well plates for 30 min in the dark, and absorbance was measured at 517 nm. For ABTS, radical cations (ABTS•⁺) were generated by reaction with potassium persulfate, diluted to an absorbance of 0.70 ± 0.10 at 734 nm, and incubated with test compounds for 6 min prior to measurement at 734 nm. In both assays, methanol or water served as negative controls, and Trolox, N-acetylcysteine, and ascorbic acid were included as positive controls. Radical scavenging activity was calculated as the percentage reduction in absorbance after correction for intrinsic compound absorbance. All measurements were performed in technical triplicates across at least three independent experiments.

### *Drosophila* lifespan and oxidative stress assays

Lifespan and oxidative stress resistance were assessed in *Drosophila melanogaster* Oregon R males. Adult flies were collected within 24 h of eclosion and maintained on standard food until day 20 post-eclosion. Flies were then transferred to food containing SR-0813 (1 or 10 μM), while control groups received food supplemented with an equivalent concentration of DMSO. For lifespan analysis, flies were housed in groups of 10 and transferred to fresh vials every 2–3 days, at which time the number of dead individuals was recorded. A total of 100 flies were analyzed per condition across four independent biological replicates. Survival was monitored until all flies had died. Survival curves were generated using the Kaplan–Meier method, and statistical significance was assessed using the log-rank (Mantel–Cox) test. For oxidative stress assays, flies were maintained on SR-0813-containing food for 10 days prior to challenge or left untreated. Flies were then starved for 1 h at 25°C in vials containing water only and subsequently transferred to vials containing 5% H₂O₂. Survival was assessed by counting the number of dead flies at 6 h and 24 h after exposure. Flies were housed in groups of 10, with a total of 100 flies per condition per replicate, and experiments were performed in three independent biological replicates. All experiments were conducted at 25°C under standard husbandry conditions.

### DHE staining of fly brains for oxidative stress imaging

Reactive oxygen species (ROS) levels in adult fly brains were assessed using dihydroethidium (DHE) staining, as previously described ^44^. Adult flies were collected at the indicated ages and briefly anesthetized on ice. Brains were dissected in cold phosphate-buffered saline (PBS) and immediately incubated with DHE (5 μM) in PBS for 15 min at room temperature in the dark. Following incubation, brains were washed three times in PBS to remove excess dye and mounted in PBS or mounting medium for imaging. Fluorescence was acquired using an Olympus FV3000 confocal microscope with a 10× objective, using identical acquisition settings across all conditions to allow quantitative comparison. Whole brains were analyzed. ROS levels were quantified by measuring fluorescence intensity using ImageJ software. In addition, a ROS index was calculated as total fluorescence intensity × (number of objects / mean number of objects in the control group), using CellSens software (Olympus). Background-subtracted mean fluorescence intensity was calculated for each brain and normalized to control samples. A total of 17 brains were analyzed per condition across three replicates.

### *Drosophila* eye degeneration modifier assays

Genetic modifier screening was performed as previously described ^20^, with minor modifications. Virgin females of the genotype *GMR-GAL4; +; dENL/AF9 IR* were crossed with males carrying *UAS-linked human neurodegenerative disease genes* (*hNDG*s) listed in Supplementary Table S1. Control crosses were performed using *GMR-GAL4; +; GFP IR* females. Crosses were maintained at 28°C, 55% humidity, under a 12 h light/dark cycle. Male F1 progeny were analyzed for suppression or enhancement of eye degeneration phenotypes relative to controls. A total of 50 males per genotype were examined across three independent parental crosses. Eye degeneration was quantified using a semi-quantitative scoring system. Flies were evaluated for loss of pigmentation, neuronal death (black spots), retinal collapse, and ommatidial fusion. Each phenotype was assigned a score based on severity: 1 point when present, increasing to 2, 3, or 4 points if the affected area exceeded 5%, 30%, or 65%, respectively. Scoring was performed independently by three investigators, and final scores were used to classify genetic interactions as suppressors, enhancers, or neutral relative to controls.

### Larval crawling (locomotor) assay

Larval locomotor activity was assessed using a crawling assay as previously described ^17^, with minor modifications. Third instar larvae were collected, briefly rinsed in distilled water to remove food residue, and placed individually on agar plates. After a short acclimation period (∼1 min), larval movement was recorded for 30 s under consistent conditions. Larval trajectories were analyzed using ImageJ (version 1.53K) with the wrMTrck plugin. Total distance traveled and path length over the 30 s interval were quantified, and average speed was calculated accordingly. Data were plotted for each condition. The number of larvae analyzed per condition is indicated in the corresponding figure legends. Larvae were obtained from three independent parental crosses. All assays were performed under standardized conditions at room temperature.

### Adult negative geotaxis (climbing) assay

Adult locomotor performance was assessed using a negative geotaxis assay as previously described ^45^, with minor modifications. Male flies were collected from the indicated genotypes and maintained under standard conditions at either 25°C or 28°C, as specified in the corresponding figures. For genetic experiments, flies expressing *UAS-UBQLN2^P^*^497^*^H^*under the control of elav-GAL4 were combined with either *dENL/AF9 RNAi* (*elav>dENL/AF9 IR>UBQLN2^P^*^497^*^H^*) or control RNAi (*elav>GFP IR>UBQLN2P497H*), and locomotor performance was assessed longitudinally. For pharmacological experiments, flies expressing neurodegenerative disease-associated transgenes (*elav>UBQLN2^P^*^497^*^H^*, *elav>SOD1^G94A^*, *elav>TDP-43^Q^*^331^*K*, or *elav>(GGGGCC)_49_*) were maintained on food supplemented with SR-0813 (0, 1, or 10 μM), with DMSO as vehicle control. At the indicated ages (5, 10, 15, 20, 25, and 30 days post-eclosion), flies were gently transferred without anesthesia into transparent conical tubes (10 flies per tube) and allowed to acclimate for 5 min. Flies were then tapped to the bottom, and climbing behavior was recorded over a 5 s interval. Each assay was repeated five times per group at 30 s intervals. Climbing performance was scored based on the height reached within 5 s using a semi-quantitative scale: 0 (<2.0 cm), 1 (2.0–3.9 cm), 2 (4.0–5.9 cm), 3 (6.0–7.9 cm), 4 (8.0–9.9 cm), and 5 (>10 cm). A climbing index was calculated as the weighted sum of scores divided by the total number of flies per group. Unless otherwise specified, 100 flies were analyzed per condition across three independent biological replicates. Performance was compared to age-matched controls.

### Neuromuscular junction (NMJ) immunostaining and morphology quantification

Third instar larvae expressing UBQLN2^P497H^ in all neurons (elav-GAL4) were treated with the indicated compounds or vehicle control, as specified in the corresponding figures. Larvae were dissected in hemolymph-like saline (HL3) and fixed in 4% paraformaldehyde in PBS for 30 min at room temperature, following established protocols. Samples were then blocked in PBS containing 2% bovine serum albumin (BSA) and 0.1% Triton X-100. For immunostaining, neuromuscular junctions were labeled using FITC-conjugated goat anti-horseradish peroxidase (HRP) antibody (1:1000, Jackson ImmunoResearch), which marks neuronal membranes, and mouse anti-Discs large (Dlg) antibody (1:300, clone 4F3, DSHB). After primary incubation, samples were washed and incubated with Alexa Fluor 594-conjugated anti-mouse secondary antibody to visualize Dlg. Preparations were mounted and imaged using an Olympus Fluoview FV3000 confocal microscope under identical acquisition settings across conditions. Quantification was performed at muscle 4 of abdominal segment A4, focusing on type Ib glutamatergic motor neuron terminals. Bouton number and NMJ length were measured using ImageJ (version 1.53k). Analyses were performed on 30 larvae derived from two independent parental crosses, with one NMJ quantified per larva per condition.

### Cell culture and neuronal differentiation

Human neuroblastoma SH-SY5Y cells were obtained from American Type Culture Collection (ATCC CRL-2266) and maintained in a 1:1 mixture of EMEM and F-12 medium supplemented with 10% fetal bovine serum and 1% penicillin–streptomycin. Cells were cultured at 37 °C in a humidified incubator with 5% CO₂ and used between passages 6–12. For neuronal differentiation, cells were seeded at 1 × 10⁵ cells per well in 6-well plates and allowed to attach for 24 h. Differentiation was initiated by switching to medium containing 10 μM retinoic acid (RA) and maintained for 3 days, with media refreshed every 48 h. Neuronal differentiation was verified by morphological changes and expression of neuronal markers. All experiments involving cell culture were approved by the Institutional Biosafety Committee (ICB) of the Faculty of Medicine, Chiang Mai University (Thailand), under protocol number CMUIBC0268022.

### Chemical treatments and stress paradigms in differentiated SH-SY5Y neurons

RA-differentiated SH-SY5Y neurons were subjected to defined chemical stressors to model distinct cellular insults. Cells were treated for 48 h with hydrogen peroxide (H₂O₂; oxidative stress), MG132 (proteasome inhibition/proteotoxic stress), tunicamycin (Tm) (endoplasmic reticulum stress), or rotenone (mitochondrial dysfunction), at concentrations indicated in figure legends. For co-treatment conditions, cells were exposed to stressors in the presence or absence of SR-0813 at the indicated concentrations. Vehicle (DMSO) controls were included in all experiments. Treatments were performed under standard culture conditions following neuronal differentiation. Cellular responses were quantified after 48 h as described in the corresponding assay sections.

### Cell viability assay (MTT)

Cell viability was assessed using the MTT assay (3-(4,5-dimethylthiazol-2-yl)-2,5-diphenyltetrazolium bromide) in RA–differentiated SH-SY5Y neurons. Cells were seeded in 96-well plates at a density of 3.3 × 10³ cells per well and differentiated for 3 days prior to treatment. Neurons were exposed to the indicated stressors (tunicamycin, H₂O₂, MG132, or rotenone) in the presence or absence of SR-0813 at concentrations and durations specified in the figure legends. Following treatment, medium was replaced with MTT solution (0.5 mg/mL) and incubated for 4 h at 37 °C. The solution was then removed, and formazan crystals were solubilized in DMSO. Absorbance was measured at 570 nm using a microplate reader. Cell viability was expressed as a percentage relative to DMSO-treated controls. Unless otherwise specified, each condition included at least six technical replicates across a minimum of two independent biological replicates.

### Measurement of intracellular ROS in SH-SY5Y cells

Intracellular reactive oxygen species (ROS) levels were measured in undifferentiated and RA–differentiated SH-SY5Y cells using the fluorescent probe (5-(and-6)-chloromethyl-2′,7′-dichlorodihydrofluorescein diacetate) (CM-H₂DCFDA). Cells were treated with Tm (1 μM) in the presence or absence of SR-0813 (0.4 or 1.6 μM), as indicated. For imaging-based analysis, undifferentiated cells were incubated with CM-H₂DCFDA, washed, and imaged by confocal microscopy using a 20× objective under identical acquisition settings. Fluorescence intensity was quantified using CellSens (Evident) software and expressed as arbitrary units (a.u.). Experiments were performed in four independent biological replicates. For flow cytometry analysis, differentiated cells were stained with CM-H₂DCFDA, harvested, and analyzed on a DxFLEX flow cytometer (Beckman Coulter, CA, USA) with excitation at 488 nm and fluorescence detected in the FITC channel. At least 10,000 events were acquired per sample. The main cell population was identified based on forward and side scatter (FSC-A and SSC-A) to exclude debris. ROS levels were quantified as the median fluorescence intensity (MFI) of DCF. Unless otherwise specified, experiments were performed with at least three independent biological replicates, and statistical significance was defined as p < 0.05.

### Apoptosis and necrosis analysis by Annexin V/PI flow cytometry

Annexin V/propidium iodide (PI) staining (FineTest, China) followed by flow cytometry was used to assess apoptosis and necrosis in RA–differentiated SH-SY5Y cells under tunicamycin (1 μM) or MG132-induced stress, in the presence or absence of SR-0813 (0.4 or 1.6 μM). Following treatment, cells were collected, washed with cold PBS, and resuspended in binding buffer. Samples were incubated with fluorophore-conjugated Annexin V and PI for 15 min at room temperature in the dark and analyzed without fixation. Cell populations were gated based on forward and side scatter (FSC-A and SSC-A) to exclude debris. Early apoptotic (Annexin V⁺/PI⁻), late apoptotic/necrotic (Annexin V⁺/PI⁺), and necrotic (Annexin V⁻/PI⁺) populations were defined based on staining profiles. Data are presented as the percentage of cells in each population, with total apoptotic cells calculated as the sum of Annexin V–positive populations. Experiments were performed with at least three independent biological replicates.

### Preparation of crude extracts and Western blotting

Crude protein extracts were prepared from RA-differentiated SH-SY5Y cells and adult *Drosophila* melanogaster heads. For SH-SY5Y cells, cultures were washed with PBS, collected, and lysed in RIPA buffer supplemented with protease inhibitors. Lysates were incubated on ice, clarified by centrifugation at 14,000 × g for 20 min at 4 °C, and the supernatant was collected. Protein concentration was determined using a BCA assay. For *Drosophila*, crude extracts were prepared by homogenizing 10 adult male heads per sample in 100 μL of 4× sample buffer. Samples were obtained from three independent biological replicates, each derived from separate parental crosses. Equal amounts of protein (30 μg for SH-SY5Y lysates) or equal volume of crude exctracts (10 μL for *Drosophila* samples) were resolved by SDS–PAGE on 12% polyacrylamide gels and transferred to PVDF membranes using a semi-dry transfer system. Membranes were blocked in 5% skim milk in TBS-T (Tris-buffered saline with 0.1% Tween-20) and incubated overnight with primary antibodies (listed in Supplementary Table S3), followed by HRP-conjugated secondary antibodies (1:3000, 1 h). Membranes were developed using WesternSure Chemiluminescent Reagents (LI-COR) and imaged with the C-DiGit Blot Scanner (LI-COR). Protein levels were normalized to housekeeping controls (e.g., Actin or Lamin C), and phosphorylated proteins were normalized to their corresponding total protein levels. Band intensities were quantified using ImageJ (v1.53k). All experiments were performed with three independent biological replicates, unless otherwise indicated.

### Quantitative RT-PCR

Total RNA was extracted from adult *Drosophila melanogaster* heads by pooling 40 heads per sample using the RNeasy Mini Kit (QIAGEN) according to the manufacturer’s instructions. Three independent biological replicates were analyzed, each derived from separate parental crosses. RNA concentration and purity were assessed spectrophotometrically. For each sample, 1 μg of total RNA was reverse transcribed into cDNA using SensiFAST cDNA Synthesis Kit (Bioline). Quantitative PCR was performed using SensiFAST SYBR® Lo-ROX Kit on a CFX Opus Real-Time PCR System (Bio-Rad), with specific primers (Supplementary Table 4). Quantitative normalization for each sample was performed by using *ACTIN* (*ACT5C*) or *GAPDH* as an internal control.

### Statistical analysis

Statistical analyses were performed using GraphPad Prism 11 (GraphPad Software). Data are presented as mean ± SD, with individual data points shown where appropriate. For comparisons between two groups, unpaired two-tailed Student’s t tests were used, with Welch’s correction applied when variances were unequal. For multiple-group comparisons, one-way ANOVA was performed followed by the multiple-comparisons test specified in the figure legend (Dunnett’s, Šidák’s, or Tukey’s). When data did not meet parametric assumptions, nonparametric tests were applied (Mann–Whitney test for two groups; Kruskal–Wallis test with Dunn’s post hoc test for multiple groups). Survival curves were analyzed using the log-rank (Mantel–Cox) test. Sample sizes, number of biological replicates, and statistical tests are indicated in the corresponding figure legends. Statistical significance was defined as p < 0.05.

